# Gene regulatory evolution in cold-adapted fly populations neutralizes plasticity and may undermine genetic canalization

**DOI:** 10.1101/2022.01.14.476421

**Authors:** Yuheng Huang, Justin B. Lack, Grant T. Hoppel, John E. Pool

**Affiliations:** Laboratory of Genetics, University of Wisconsin-Madison, Madison, Wisconsin 53706

**Author notes:** Current address: Department of Ecology and Evolutionary Biology, University of California, Irvine, Irvine, CA 92697. Current address: Advanced Biomedical Computational Science, Frederick National Laboratory for Cancer Research, Frederick, MD 21701.

**Keywords:** adaptive evolution, transcriptomic plasticity, genetic canalization, *Drosophila melanogaster*

## Abstract

The relationships between adaptive evolution, phenotypic plasticity, and canalization remain incompletely understood. Theoretical and empirical studies have made conflicting arguments on whether adaptive evolution may enhance or oppose the plastic response. Gene regulatory traits offer excellent potential to study the relationship between plasticity and adaptation, and they can now be studied at the transcriptomic level. Here we take advantage of three closely-related pairs of natural populations of *Drosophila melanogaster* from contrasting thermal environments that reflect three separate instances of cold tolerance evolution. We measure the transcriptome-wide plasticity in gene expression levels and alternative splicing (intron usage) between warm and cold laboratory environments. We find that suspected adaptive changes in both gene expression and alternative splicing tend to neutralize the ancestral plastic response. Further, we investigate the hypothesis that adaptive evolution can lead to decanalization of selected gene regulatory traits. We find strong evidence that suspected adaptive gene expression (but not splicing) changes in cold-adapted populations are more vulnerable to the genetic perturbation of inbreeding than putatively neutral changes. We find some evidence that these patterns may reflect a loss of genetic canalization accompanying adaptation, although other processes including hitchhiking recessive deleterious variants may contribute as well. Our findings augment our understanding of genetic and environmental effects on gene regulation in the context of adaptive evolution.

**Significance Statement:** It is unclear whether adaptive evolution is concordant or discordant with regulatory plasticity, especially for splicing plasticity which is rarely studied. Here we analyzed RNA-seq data from three pairs of natural fly populations that represent separate adaptive evolution to cold climate. We found that adaptive evolution is generally discordant with the ancestral plasticity between cold and warm temperatures for gene expression abundance and splicing. We also investigate the hypothesis that adaptation leads to decanalization of the selected traits. By comparing the expression variance between inbred and outbred samples, we found evidence that adaptation may lead to genetic decanalization for expression abundance but not for splicing. Our study reveals the relationship between adaptation, plasticity and canalization in three instances in nature.

## Introduction

For organisms to cope with environmental changes, two important strategies are adaptive evolution and phenotypic plasticity (Meyers & Bull, 2002). Phenotypic plasticity is the phenomenon of a single genotype producing different phenotypes under different environmental conditions. Producing different phenotypes from a genotype often requires an intermediate step, such as gene expression change (Ghalambor et al., 2007; Pfennig et al., 2010). It has also been shown in many cases that expression evolution contributes to adaptive evolution (e.g., Fraser et al., 2010; Nourmohammad et al., 2017). Studying the interactions between expression plasticity and adaptive expression evolution can generate insights on how these two processes help organisms respond to environmental changes.

Adaptation to a new environment may change gene expression in the same direction as the initial plasticity, which suggests the initial plasticity is beneficial, shifting the phenotype toward the optimal phenotypic values in the selective environment (Chevin et al., 2010; Via, 1993). Alternatively, a plastic response can be deleterious if it shifts the phenotype away from the optimum under environmental perturbation and adaptive evolution should restore the phenotype toward the ancestral state (Scheiner, 1993; von Heckel et al., 2016). The latter scenario predicts a negative relationship between the direction of plastic change and that of evolutionary change, resulting in a “counter-gradient” pattern (Conover & Schultz, 1995). Finally, environmentally induced expression change may have little effect on fitness. In that case, evolutionary change is determined by drift, showing no relation with the plasticity in terms of direction. Many recent laboratory and field studies have observed a “counter-gradient” pattern for plasticity and adaptive evolution for gene expression (e.g., Dayan et al., 2015; Fischer et al., 2021; Ghalambor et al., 2015; Ho & Zhang, 2018; Huang & Agrawal, 2016; Koch & Guillaume 2020; Levine et al., 2011; Ragland et al., 2015) while some other studies found that adaptive plasticity is more common (Bittner et al., 2021; Josephs et al., 2021; Kenkel & Matz, 2017; Mallard et al., 2020).

However, few of them have studied multiple natural population pairs that reflect separate adaptation events. Moreover, the relationship between plasticity and adaptive evolution for other regulatory traits such as splicing is unexplored.

On the other hand, adaptation to a new environment might have a significant consequence on the level of canalization of the selected trait. Canalization refers to the capacity of a trait to maintain constant phenotype under genetic or environmental perturbation (Waddington 1942; Flatt, 2005). This concept has been defined operationally in different ways (Dworkin 2005). Environmental canalization has been described as the opposite of phenotype plasticity (Nijhout and Davidowitz 2003), as indicated by the constancy of a trait across environments. Hence, plasticity can be viewed as a type of environmental de-canalization. However, environmental canalization can alternatively be defined based on the trait variation observed among individuals in one environment compared to others (Stearns and Kawecki 1994). Similarly, genetic canalization can be described as the constancy of the phenotype under heritable perturbations, such as mutations (de Visser et al. 2003). It can be measured as the inverse of the variance caused by mutations (Flatt, 2005).

Canalization can potentially constrain evolution by reducing phenotypic variation (Charlesworth et al. 1982; Maynard Smith et al. 1985), or it may facilitate evolution by allowing cryptic genetic variation to accumulate (Gibson & Dworkin, 2004; Masel, 2005; Paaby & Rockman, 2014). If a trait is under stabilizing selection in the ancestral environment, its developmental canalization may be favoured by selection (Gavrilets & Hastings, 1994; Wagner et al., 1997). However, when adaptive evolution shifts a trait to a new optimum, the previous canalization mechanism may be undermined. This possibility, that adaptive evolution could result in decanalization, was hinted at when blowflies with newly-evolved insecticide resistance were found to have bristle asymmetry and prolonged development (Clarke & McKenzie, 1987; McKenzie & Game, 1987), and also when in vitro selection on an enzyme resulted in reduced robustness to both genetic and environmental perturbations (Hayden et al. 2012). A natural example of decanalization of the same trait that evolved adaptively was then provided by Lack et al. (2016b), who found that highland Ethiopian *D. melanogaster* with derived large wing size had greatly reduced genetic robustness of wing development. One subsequent study found that environmental decanalization may not have occurred for these Ethiopian fly wings (Pesevski & Dworkin 2020), highlighting the potential distinctness of genetic and environmental canalization. A further study showed that when wing size in the same species was artificially selected, both genetic and environmental decanalization of wing development ensued (Groth et al. 2018). Whereas, longer-term lab selection of *D. melanogaster* for rapid development was suggested to increase canalization of that trait (Ghosh et al. 2019). Whether a decanalizing effect of directional selection exists for a substantial proportion of adaptive gene regulatory traits has not previously been investigated, and this question represents one focus of the present study.

In this study, we take advantage of African and European populations of *Drosophila melanogaster* that have experienced parallel adaptation to colder climates, to study the interaction between adaptation and plasticity as well as canalization in gene expression. Originating from a warm sub-Saharan range (Lachaise et al., 1988; Pool et al., 2012; Sprengelmeyer et al., 2020), populations have independently occupied colder environments at least three times: in higher latitudes of Eurasia (here represented by the France FR population), and in the highlands of Ethiopia (EF population) and South Africa (SD population). Flies from these three locations were paired with genetically similar populations from warm regions: Egypt (EG), Ethiopia lowland (EA), and South Africa lowland (SP), respectively, together representing three different population pairs: Mediterranean (MED), Ethiopian (ETH), and South African (SAF). These cold-adapted populations have evolved separately from the warm-adapted ones for ~1,000-2,000 years (~15,000-30,000 generations) (Sprengelmeyer et al., 2020). They show parallel changes in cold tolerance, as measured by recovery after prolonged 4°C cold exposure (Pool et al., 2017) and greater egg-to-adult survival at 15°C (Huang et al., 2021). These cold-adapted populations also show a genome-wide excess of parallel allele frequency shifts (Pool et al. 2017).

In our related study (Huang et al. 2021), we identified evolutionary changes in gene expression abundance and alternative splicing (intron usage) between cold- and warm-adapted populations for each pair at a low temperature, approximating the derived cold environment. Here, we rear the same crosses from these populations at a warm temperature, approximating the ancestral environment, and analyze transcriptomes of adult females. First, we compare the directions of gene regulatory plasticity between cold and warm environments and the directions of evolutionary changes between cold and warm populations for both expression abundance and intron usage. We find adaptive evolution tends to counter plasticity by restoring the ancestral regulation (neutralizing the ancestral plasticity). Second, since we previously analyzed gene regulation from some parental inbred lines of the outbred crosses we otherwise analyze (Huang et al. 2021), we use inbreeding as a form of genetic perturbation (Réaale & Roff, 2003; de Visser et al., 2003) to study regulatory canalization in the cold and warm populations and its potential disruption due to adaptive evolution. We use the ratio of regulatory trait variance among inbred samples to that among outbred samples as a potential signal of genetic decanalization. We indeed find transcriptome-wide evidence consistent with adaptive evolution leading to decanalization for expression abundance traits, although other processes such as hitchhiking deleterious variants many contribute to this signal as well.

## Results

### Adaptive evolution and naïve plasticity in expression and splicing

We collected RNA-seq data from whole female adult samples representing the three warm/cold population pairs described above, raised at temperatures intended to reflect either an ancestral warmer environment (25°C) or a derived cooler environment (15°C). For each of these six populations, the transcriptomes from eight unique outbred crosses were analyzed. We first performed PCA on the normalized expression read counts across samples from different environments and populations. There are signals of temperature environment on transcriptome variation, separating the samples from the cold environment from the respective samples from the warm environment along the PC1 (Fig. S1). Then we characterized the genes/introns that showed consistent plastic expression/splicing changes between 15°C and 25°C rearing environments in warm-adapted populations (*P*). Formally, we required the direction of plasticity for at least seven out of the eight crosses to be consistent with the average plastic change (Materials and Methods). We refer to this pattern from warm-adapted populations as ancestral or naïve plasticity, hypothetically representing the initial influence of derived cooler environments on expression/splicing prior to any cold adaptation. Across the six populations (three warm/cold-adapted population pairs), 14-65% of all gene expression traits met the consistent plasticity criterion (at least seven out of eight crosses showing the same direction of plastic change), as did 24-36% of intron usage traits (Table 1A). The Mediterranean pair had the greatest proportions of consistently plastic traits for both expression and splicing. Across all populations and traits, an average 29.9% genes/introns show consistent plasticity by our criteria. In light of our expected false positive rate (7%), we estimate that approximately 77% of the identified genes are true positives for plasticity, which provides a substantially enriched set of plasticity candidates for the transcriptome-wide analyses described below.

**Table 1.**
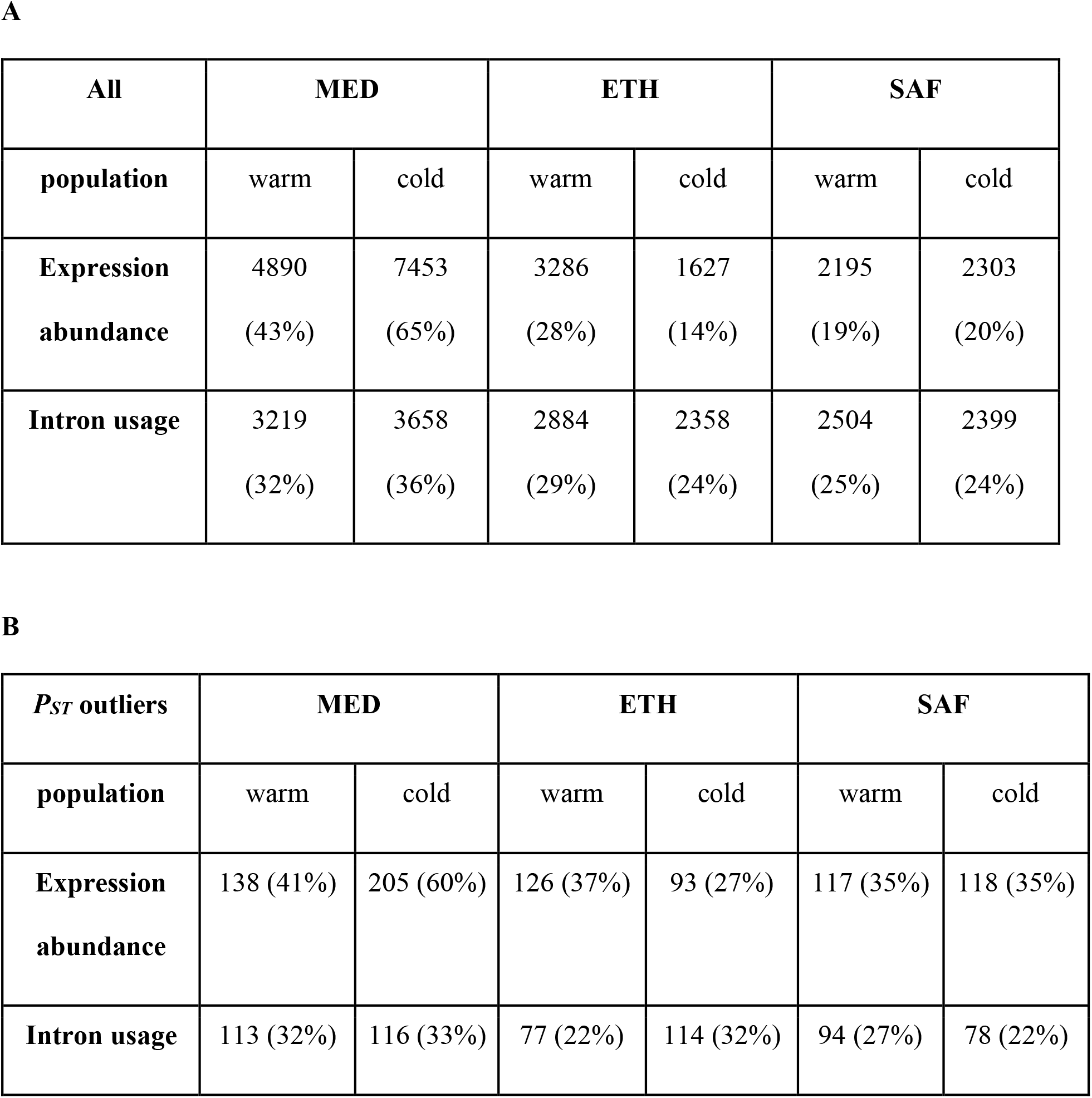
Numbers of gene expression abundance and intron usage traits showing plasticity in warm- and cold-adapted populations for each population pair. Panel A shows all the genes/introns and panel B shows those with elevated expression/splicing differentiation between adult females from cold- and warm-adapted populations at 15°C (*P_ST_* outliers identified in Huang et al. 2021). The percentage in parentheses indicates the proportion showing consistent plasticity.

| **All** | **MED** |  | **ETH** |  | **SAF** |  |
| --- | --- | --- | --- | --- | --- | --- |
| **population** | warm | cold | warm | cold | warm | cold |
| **Expression abundance** | 4890<br>(43%) | 7453<br>(65%) | 3286<br>(28%) | 1627<br>(14%) | 2195<br>(19%) | 2303<br>(20%) |
| **Intron usage** | 3219<br>(32%) | 3658<br>(36%) | 2884<br>(29%) | 2358<br>(24%) | 2504<br>(25%) | 2399<br>(24%) |

Table 1. Numbers of gene expression abundance and intron usage traits showing plasticity in warm- and cold-adapted populations for each population pair.
| ***P<sub>ST</sub>* outliers** | **MED** |  | **ETH** |  | **SAF** |  |
| --- | --- | --- | --- | --- | --- | --- |
| **population** | warm | cold | warm | cold | warm | cold |
| **Expression abundance** | 138 (41%) | 205 (60%) | 126 (37%) | 93 (27%) | 117 (35%) | 118 (35%) |
| **Intron usage** | 113 (32%) | 116 (33%) | 77 (22%) | 114 (32%) | 94 (27%) | 78 (22%) |

We then set out to assess the general relationship between a warm-adapted population’s naïve plasticity and the evolved difference between the populations in the derived environment. Here, we are asking whether evolutionary changes are concordant or discordant with naïve plasticity, as opposed to asking how plasticity itself has evolved. We compared the directions of initial plasticity (*P*) in warm-adapted populations and evolved change (*E*), which is the difference in expression between cold and warm-adapted populations at 15°C. If *P* and *E* were in the same direction (i.e., *P* ◻ *E* > 0), the evolved change was defined as “concordant” (Fig. 2). Considering all plastic genes for gene expression abundance, the proportion of “concordant” changes was 53% in MED, 55% in ETH, and 39% in SAF. For intron usage, the proportion of “concordant” changes was 41% in MED, 53% in ETH, and 38% in SAF.

**Fig. 1.**
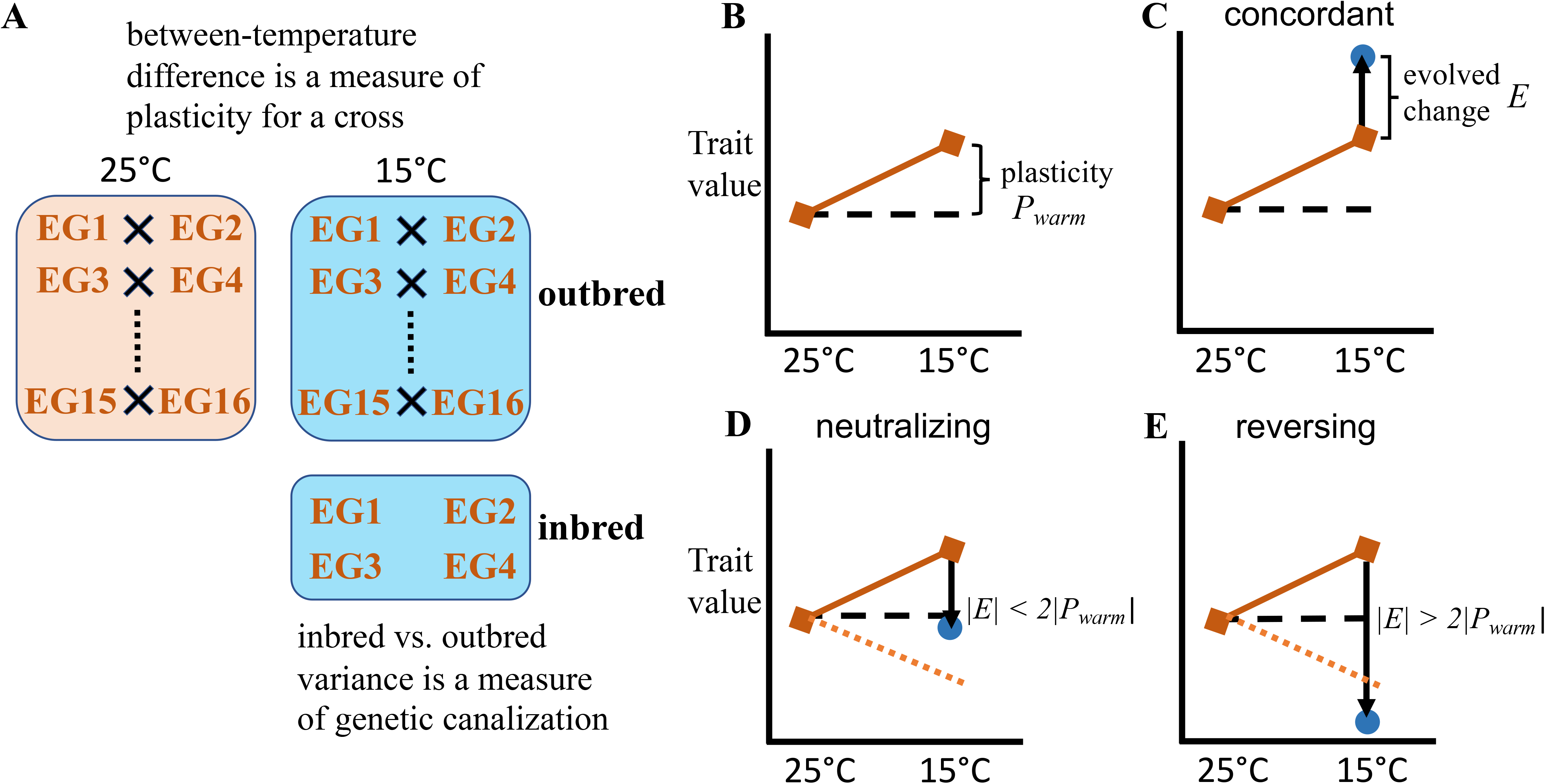
Illustrations of the experimental design and different expectations for adaptative changes regarding the naïve plasticity. (A) Experimental design is shown for one population (EG, a warm population) as an example. Crosses generated outbred offspring at 15°C (derived condition) and 25°C (ancestral condition) to study transcriptome plasticity. Inbred strains were reared at 15°C to study genetic canalization by comparing the variance of trait value for inbred vs. outbred samples at 15°C. Whole female adults were collected from the offspring. (B-E) Conceptual illustrations of different expectations for evolved changes and naïve plasticity. (B) Naïve plasticity for a warm population. The difference between trait values at 15°C vs. 25°C is a measure of plasticity (*P_warm_*). (C) The evolved change (*E*, solid arrow) is the difference between trait values between cold (blue dot) and warm population (orange square) at 15°C. If the evolved change is in the same direction as the naïve plasticity, the evolved change is regarded as “concordant”. (D) If the evolved change is in the opposite direction as the naïve plasticity (*E* ⍰ *P_warm_* < 0) and moves the trait closer to the warm population’s trait value at 25°C (|*E*| < 2|*P_warm_*|), it is regarded as “neutralizing”. The orange dashed line indicates the level of plastic change but in the opposite direction. (E) If the evolved change moves the trait further from the warm population’s trait value at 25°C with a magnitude greater than the initial plasticity in the opposite direction (*E* ⍰ *P_warm_* < 0; |*E*| < 2|*P_warm_*|), it is regarded as “reversing”. The “concordant” and “reversing” categories are both regarded as “novelty” for the canalization analysis, because they each take the cold-adapted population’s trait value in its home environment farther away from the warm-adapted population’s trait value in its own home environment than naïve plasticity would have done on its own.

**Fig. 2.**
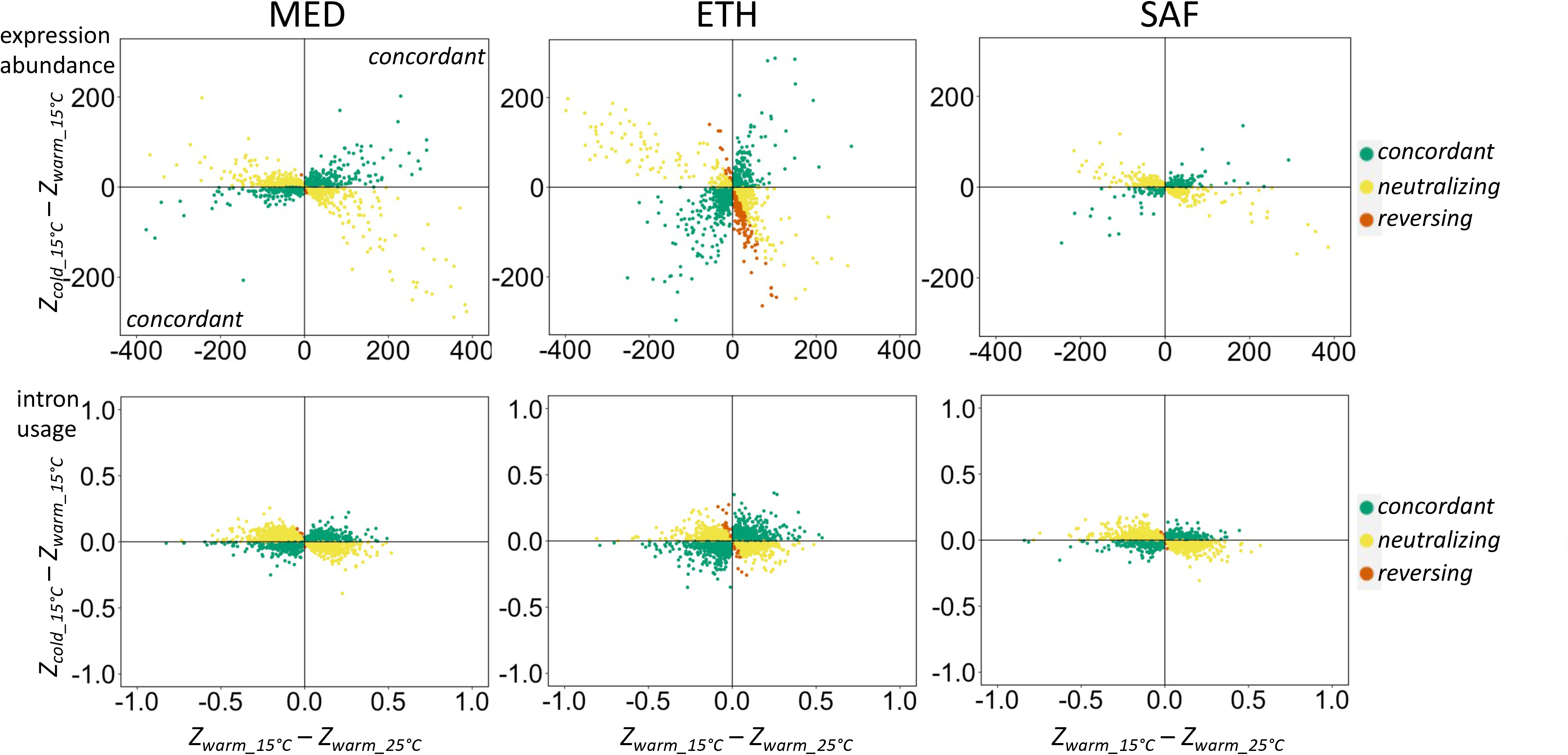
The plastic responses of the warm-adapted population (*P_warm_*, x-axis) and the evolved differences in expression between populations in the derived cold environment (*E*, y-axis) for all genes (upper panel) and introns (lower panel) that showed consistent naïve plasticity. Dots located in the lower left and upper right quadrants indicate that the plasticity responses and evolutionary changes are concordant.

We further partitioned those traits where *P* and *E* were in opposite directions (“discordant”, *P* ◻ *E* < 0). If the evolved change moved expression/splicing farther from ancestral levels than the ancestral plasticity did, (the magnitude of evolved change was more than twice that of plasticity, |*E*| > |2*P|*), we defined it as “reversing”. If instead, the evolved change brought the cold population at 15°C back closer to the warm population at 25°C, then we defined the evolved change as “neutralizing”. These categories may have distinct evolutionary interpretations, since neutralizing changes are consistent with a single trait optimum across thermal environments, whereas for reversing changes, the observation that evolution increased trait differentiation between the populations in their home environments could indicate distinct adaptive optima in cold vs. warm environments. We found that among “discordant” changes, “reversing” was a relatively small fraction across population pairs (Fig. 2; on average 11% for expression abundance and 1.7% for intron usage). It is worth noting that the Ethiopia pair appears to have a much higher proportion of “reversing” changes (32% for expression and 3.6% for intron usage) than the other two pairs, suggesting distinct evolution in the Ethiopia pair.

To examine whether putative adaptive evolution in expression/splicing in cold populations followed the initial plastic changes, we focused on genes showing elevated population differentiation in gene expression abundance or intron usage between warm- and cold-adapted populations based on Huang et al. 2021 (about 339 outlier genes and 351 outlier intron junctions for each population pair). These candidates for adaptive gene regulatory evolution were identified using a *P_ST_* outlier approach, focusing on the top 5% of genes/introns for each population pair separately (Huang et al. 2021; Materials and Methods). Among these outliers (an average of 339 genes per population pair for expression and 351 introns for splicing), the numbers of expression abundance genes also passing the cutoff for plasticity were in the range of 93 to 205 across population pairs; the numbers of divergent intron usage traits passing the plasticity cutoff were in the range of 77 to 116 (Table 1B).

For both expression abundance and intron usage, we observed a general pattern that plastic *P_ST_* outliers had lower proportions of “concordant” naïve expression plasticity than plastic non-outliers across all population pairs (Fig. 3). In other words, putatively adaptively evolved divergence tended to oppose the naïve plasticity. For expression abundance, we found the difference in the “concordant” proportion was significant for the ETH (χ = 8.3, df = 1, p = 0.0039) and SAF pairs (χ^2^ = 15.1, df = 1, p = 9.9e-05) but only marginally significant for the MED pair (χ^2^ = 3.6, df = 1, p = 0.058). For intron usage, the difference in the “concordant” proportion was significant for the MED (χ^2^ = 6.7, df = 1, p = 0.0096) and the SAF pair (χ^2^ = 21.7, df = 1, p = 3.2e-06) but not the ETH pair (χ^2^ = 0.24, df = 1, p = 0.62). Mirroring transcriptome-wide patterns, a large majority of *P_ST_* outliers with “discordant” changes were “neutralizing” rather than “reversing”; the average proportion of “reversing” is 4% for gene expression and 9% for intron usage. Hence, the largest share of putatively adaptive regulatory changes served to mitigate ancestral plasticity.

**Fig. 3.**
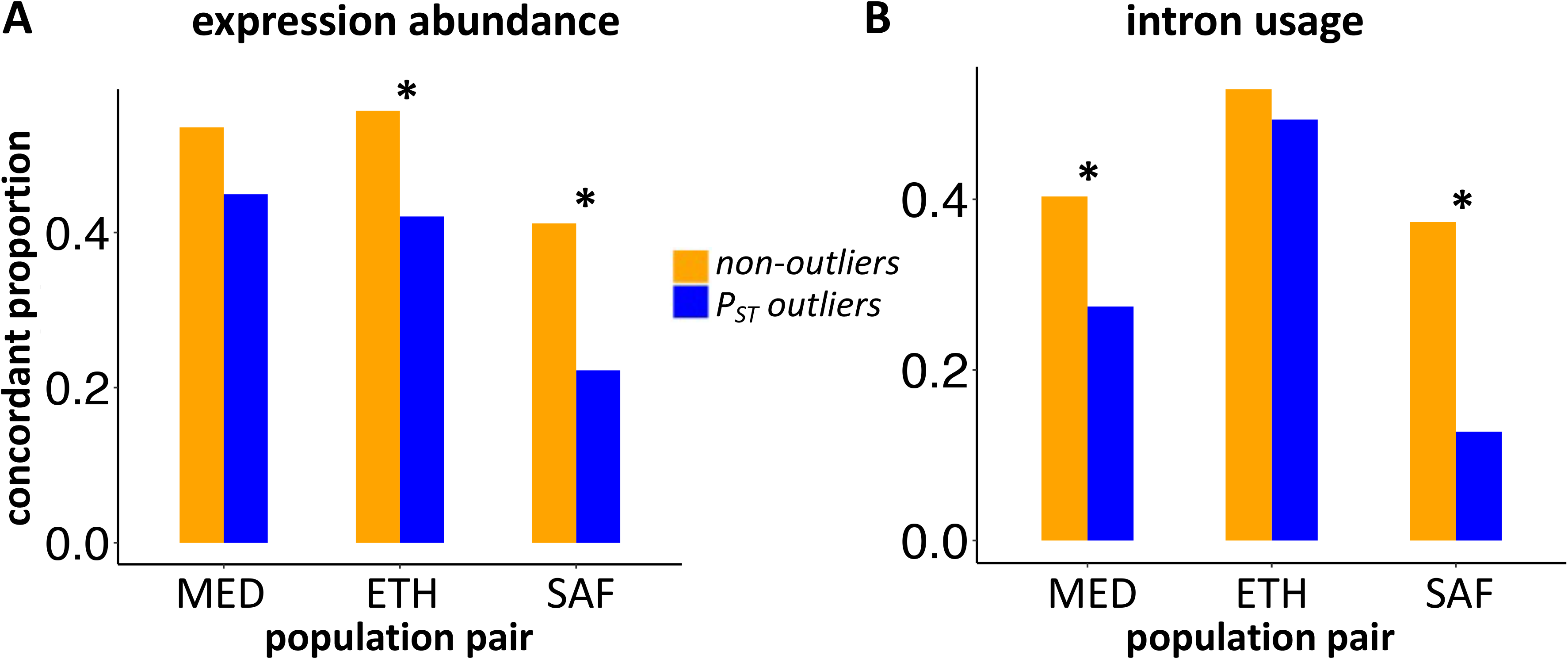
The proportion of genes/introns classified as “concordant” of naïve plasticity for *P_ST_* outliers and non-outliers. * indicates the proportion is significantly different between *P_ST_* outliers and non-outliers (p < 0.01).

Because the same data from the warm-adapted population in the cold environment was used to estimate both plastic and evolutionary changes, it is possible that random deviations in this estimate could yield a bias (Mallard et al., 2018; Ho & Zhang, 2019). For example, if this quantity was underestimated, then plasticity would become more negative but evolution would become more positive. This potential bias toward discordant changes should be more pronounced for non-outliers in light of their more uncertain direction of evolution (whereas outliers tend to have greater magnitudes of evolution, for which the direction is less likely to be altered by measurement error). This suggestion assumes that outliers reflect true evolution (adaptive or otherwise) and were not generated via measurement error; we note that each population’s extreme high and low values were excluded from *P_ST_* calculations as a strategy to reduce the influence of measurement error (Huang et al., 2021). In contrast to the above prediction based on bias, we observed above that *P_ST_* outliers had more discordant changes than non-outliers (Figure 3).

The potential bias toward discordant changes should only exist if overlapping data are used to estimate plasticity and evolution. Therefore, we repeated the above analysis using non-overlapping subsets of the data to estimate plasticity and evolution. We randomly split the shared data (the warm-adapted population in 15°C, *Z_warm_15°C_*), with four samples being used to estimate plasticity (*P_warm_* = *Z_warm_15°C_’* – *Z_warm_25°C_*) and the other four being used to estimate evolution (*E* = *Z_cold_15°C_* – *Z_warm_15°C_’*) and identify *P_ST_* outliers, for all possible subdivisions of the data. The results were qualitatively similar to those reported above (Figure S3 versus Figure 3). All three population pairs again showed an excess of discordant changes among *P_ST_* outliers for expression. For splicing, the MED and SAF pair showed a significant excess of discordant changes among outliers (Fig. S3).

### GO functional enrichment for plastic genes/splicing

Functional analysis (GO enrichment testing) of all plastic genes (regardless of *P_ST_* outlier status) with expression/splicing showing either concordant, neutralizing or reversing evolved changes is shown in Table S4. For gene expression, we found more significant GO categories for genes showing “neutralizing” changes among three population pairs, compared to “concordant” or “reversing”. While for splicing, most of the significant GO categories were found for those showing “concordant” in the MED pair (Table S4). Further, we considered the candidates for adaptive differentiation between warm- and cold-adapted populations (Huang et al. 2021) and performed similar GO enrichment tests. Power for these analyses may be reduced because only 71 – 203 adaptation candidates with consistent plasticity were assigned among our three categories (see above). We ran separate GO enrichment analyses based on whether the potential adaptive evolution is “concordant” or else “neutralizing” the initial plasticity (there were fewer than 10 genes/introns showing “reversing” changes and therefore this category was not considered). We only found significant GO terms for the MED pair. For “concordant”, there is one significant term, “lipid particle”; For “neutralizing”, the significant terms are “coenzyme binding”, “cofactor binding”, “ion transmembrane transporter activity”, “magnesium ion binding”, “NAD binding” and “transferase activity, transferring acyl groups”. Interestingly, “lipid particle” is also found in the GO terms for *P_ST_* outliers at the adult stage in MED pair (Huang et al., 2021).

### Gene regulatory evolution and genetic canalization

We then explored how selection history may influence gene regulatory canalization. Our primary interest was to test whether transcriptomic evolution supported the hypothesis that adaptive evolution may be accompanied by a loss of canalization, which was suggested in the case of *D. melanogaster* wing size evolution in nature and in the laboratory (Lack et al., 2016; Groth et al., 2018). Here, we leveraged previously-collected RNA-seq data (Huang et al., 2021) from a subset of the parental inbred strains of the outbred crosses that all other data and analyses were derived from. Inbreeding reflects a broad genetic perturbation due to recessive deleterious variants being made homozygous. Comparisons between inbred and outbred data can therefore illuminate the relative strength of genetic canalization between different groups (Réaale & Roff, 2003; de Visser et al., 2003). For example, we would predict that for a gene expression abundance trait, a population with relatively lower canalization for this gene would have greater deviations in expression level in the inbred samples compared to the outbred samples, whereas a population with greater canalization for this gene would have relatively less difference in the variability of inbred and outbred samples (less effect of the perturbation). We therefore calculated, for each relevant regulatory trait in each population, the variance of this trait among inbred (*V_inbred_*) and outbred (*V_outbred_*) samples. Inbred data was available, and hence these comparisons were made, for the colder 15°C environment only.

Although we had a particular interest in decanalization involving potential adaptive changes showing regulatory novelty, we began by comparing *V_inbred_* and *V_outbred_* across all expression and splicing traits, many of which may be evolving neutrally. For each population, the *V_inbred_* and *V_outbred_* are highly correlated (Fig S4). Compared between populations, warm-adapted populations had much lower average *V_outbred_* than cold-adapted populations (Table 2), in spite of having somewhat higher genetic diversity than their cold-adapted counterparts (Lack et al., 2016a). The reasons for this pattern are unclear; it is possible that the stressed transcriptomic profiles of warm-adapted flies made them relatively more similar. In contrast, warm-adapted populations had higher *V_inbred_* than cold-adapted populations, except the SAF pair (Table 2). Concordantly, we found that warm-adapted populations consistently had more genes and introns showing *V_inbred_* > *V*_outbred_ than cold-adapted populations (Fig. 4A&B). The much greater increase in *V_inbred_* relative to *V_outbred_* in warm-adapted populations may reflect the relatively greater stress experienced by these flies at 15°C, compared with those from populations better adapted to such conditions (in other words, the effects of genetic and environmental perturbation may be synergistic). Alternatively, the warm-adapted populations may be expected to hold a somewhat greater number of recessive deleterious variants to be exposed by inbreeding (see Discussion).

**Table 2.** Transcriptome-wide mean variance of expression or intron usage among inbred strains or outbred crosses within a population, averaged across all genes/intron usages.

|  | pair | MED |  | ETH |  | SAF |  |
| --- | --- | --- | --- | --- | --- | --- | --- |
|  | population | warm | cold | warm | cold | warm | cold |
| Expression<br>abundance | *V<sub>inbred</sub>* | 23438 | 15965 | 61003 | 37384 | 12847 | 34075 |
|  | *V<sub>outbred</sub>* | 5132 | 10412 | 5083 | 25574 | 3125 | 61581 |
| Intron<br>usage | *V<sub>inbred</sub>* | 0.0071 | 0.0067 | 0.0113 | 0.0095 | 0.0064 | 0.0073 |
|  | *V<sub>outbred</sub>* | 0.0039 | 0.0059 | 0.0033 | 0.0049 | 0.0042 | 0.0089 |

**Fig. 4.**
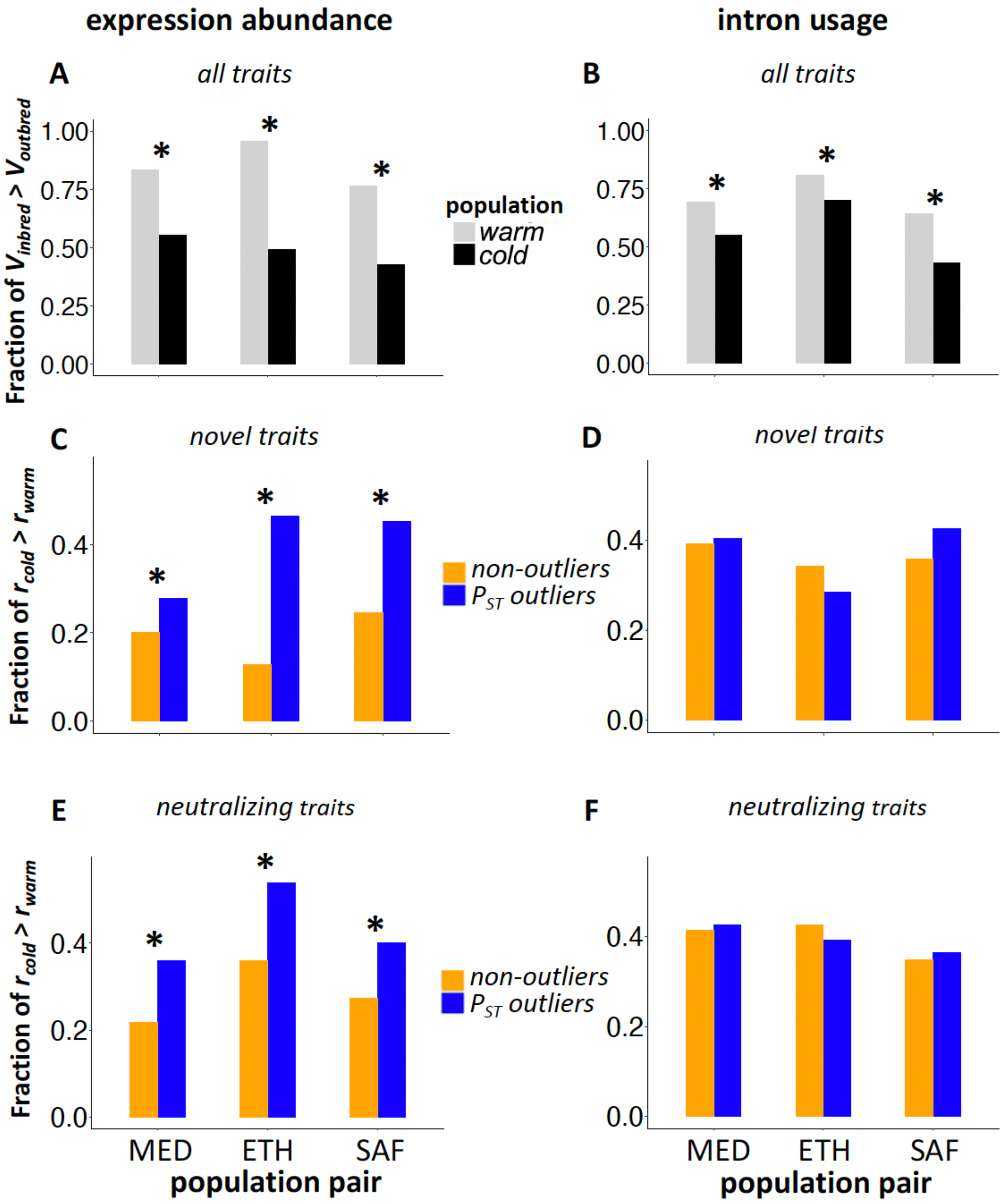
Genetic decanalization measured by *V_inbred_* relative to *V_outbred_* (*r*). A & B: The transcriptome-wide fraction of regulatory traits for which *r* > 1 for each warm-adapted and cold-adapted population is shown for expression abundance (A) and intron usage (B). C – F: The fraction of traits with novel regulatory changes (C & D) or with neutralizing changes (E & F) for which *r_cold_* > *r_warm_* for *P_ST_* outliers and non-outlier controls is shown for expression abundance (left) and intron usage (right). * indicates that the fractions are significantly different (all with p < 0.05).

Regardless of the forces shaping transcriptome-wide patterns of inbred and outbred variance, we were interested in asking whether genes potentially under recent adaptive regulatory evolution (*P_ST_* outliers) show distinct patterns of inbred versus outbred trait variance consistent with decanalization, compared to other genes. If adaptive evolution has resulted in decanalization, we predict that *P_ST_* outliers will have elevated ratios of *V_inbred_* to *V_outbred_* in cold-adapted populations compared with their warm-adapted counterparts. However, one alternative explanation for such a result is the hitchhiking of deleterious recessive regulatory variants, since any given selected haplotype may have linked harmful variants that increase in frequency alongside the favored variant (e.g. Chun & Fay 2011; Good & Desai 2014). Fortunately, these competing hypotheses make contrasting predictions with regard to adaptive changes that produce “novel” states distinct from those observed in ancestral range populations versus those “neutralizing” naïve plasticity to maintain ancestral regulatory states. The decanalization hypothesis hinges on adaptation producing new phenotypes that are not fully buffered by ancestral canalization mechanisms. Therefore, this hypothesis would predict that novel adaptive regulatory changes should have greater potential to undermine ancestral buffering mechanisms than neutralizing changes, and therefore novel outliers should have greater ratios of inbred versus outbred variance. Whereas, the deleterious hypothesis depends only upon the hitchhiking process itself, and not the magnitude or direction of the adaptive regulatory change, and therefore it predicts no difference between novel and neutralizing changes.

In testing the above predictions, we first classified all expression and splicing traits based on whether their evolution was novel or “neutralizing” in the cold environment. Similar to the plasticity section, we used the two values *P_warm_* and *E* to characterize each regulatory trait (Fig. S2). To classify novel changes, we required that the evolved change took the expression value out of the ancestral plasticity range, which included both cases where *P* ⍰ *E* > 0 (“concordant”) and those where *P* ⍰ *E* < 0 and |*E*| > |2*P|* (“reversing”). The remaining traits showing *P_warm_* ⍰ *E* < 0 and |*E*| < |2*P*| were again classified as neutralizing. Averaged across three population pairs, 37% of all expression abundance and 45% of all intron usage traits were deemed novel.

We then compared novel/neutralizing results from all genes to those from *P_ST_* outliers. For each of our *P_ST_* outlier genes showing elevated differentiation between a warm and cold population (Huang et al., 2021) that also showed evidence for regulatory novelty, we therefore tested whether the *V_inbred_* / *V*_outbred_ ratio (*r*) was higher in the cold population (*r_cold_*) or the warm population (*r_warm_*). If directional selection has had important decanalizing effects on the evolved regulatory traits, we would predict that novel *P_ST_* outliers would be more likely to have a larger *r_cold_* than *r_warm_*, compared with non-outlier genes classified as novel. For gene expression abundance, we found that the fraction of genes with *r_cold_* > *r_warm_* was significantly higher for outliers than the nonoutliers for all three population pairs (Fig. 4C. MED pair: 29% vs 20%, χ^2^ = 6.7, df = 1, p = 0.0098; ETH pair: 47% vs 13%, χ^2^ = 182, df = 1, p < 2.2e-16; SAF pair: 45% vs 24%, χ^2^ = 33, df = 1, p = 8.0e-9.). The result is consistent with the hypothesis that directional selection on expression abundance led to decanalization on these same traits.

Alternatively, the above difference between novel outliers and novel non-outliers is also compatible with the hitchhiking of deleterious variants. Furthermore, a qualitatively similar excess of *r_cold_* > *r_warm_* genes was observed for neutralizing expression changes as well (Fig. 4E. MED pair: 36% vs 22%, χ^2^ = 12, df = 1, p = 0.0005; ETH pair: 54% vs 36%, χ^2^ = 13, df = 1, p = 0.0003; SAF pair: 40% vs 27%, χ^2^ = 11, df = 1, p = 0.0008.). Since the deleterious model predicts a greater probability of *r_cold_* > *r_warm_* for all types of outliers, this result for neutralizing changes might indicate a role for this hitchhiking process.

Since the decanalization model does make a stronger prediction of inbreeding disruption for novel outliers, and the deleterious model does not, we conducted a more formal test asking if novel outliers showed a more enhanced *r_cold_* > *r_warm_* probability (compared to novel non-outliers) than neutralizing outliers (compared to neutralizing non-outliers). We performed a regression GLM analysis to model the above probability as a function of outlier status, novel/neutralizing category, and we tested for a significant interaction between those two variables. The MED and SAF returned non-significant results, with different directions as indicated by Figure 4. However, the ETH pair yielded a highly significant interaction indicating a greater decanalization signal for novel than neutralizing outliers (p = 2.6e-5). In addition, the outlier enrichment of *r_cold_* > *r_warm_* was stronger for genes where neutralizing evolutionary changes “overcompensated” for plasticity (|*P*| < |*E*| < |2*P*|) versus genes where evolutionary changes “undercompensated” for plasticity (|*E*| < |*P*|; Fig. S5), which is not predicted by the deleterious model. Based on a similar regression GLM analysis as described above, this excess was significant for the MED pair specifically (Fig. S5; p = 0.0053). Under the decanalization hypothesis, it is possible that evolved expression changes that do not generate substantially novel states might still disrupt ancestral buffering mechanisms, particularly and seem more likely to do so if their magnitude of change is greater. Hence, we find some support for the decanalization hypothesis specifically.

In contrast to the results for gene expression, for intron usage, the fraction of genes with *r_cold_* > *r_warm_* was not significantly different between outliers and non-outliers for any population pair or partition of outlier introns. Two pairs showed a qualitatively similar pattern for novel changes (Fig. 4D) but just one did so for neutralizing changes (Fig. 4F). Hence, there was no clear support for decanalization or deleterious hitchhiking for the putatively adaptive alternative splicing traits examined.

## Discussion

Gene regulation can play a key role in mediating complex interactions between genotype, environment, and fitness. In this study, we use parallel cold-adapted natural fly populations to investigate the interplay between gene regulatory evolution, plastic effects of the environment on a trait, and canalization which may buffer against such effects. Regulatory plasticity may be beneficial if it is aligned with a trait’s fitness-environment gradient (*e.g.* if a gene contributing to cold tolerance is upregulated under cold conditions). In other cases, environmental stresses may trigger unfavorable shifts in gene regulation.

Here, focusing on outliers that may reflect adaptive regulatory evolution, we observe evidence of adaptive gene expression divergence opposing the initial plasticity for gene expression and splicing across three pairs of natural populations that evolved in parallel (Fig. 3). These patterns are consistent with several previous studies, which found similar “counter-gradient” patterns in expression (Levine et al., 2010; Dayan et al., 2015; Ghalambor et al., 2015; Huang & Agrawal, 2016; Ho & Zhang, 2019; Koch & Guillaume 2020). Unlike past studies, we also examine splicing evolution and find that it shows a similar pattern, suggesting this “counter-gradient” result may be general for transcriptomic traits. Further, among the expression/splicing traits showing evolution opposing the initial plasticity, the majority of them are neutralizing the initial plastic response, i.e., restoring the expression toward the ancestral state, rather than reversing changes generating novel regulatory states.

There are at least two potential reasons why adaptive evolution may neutralize the initial plasticity. First, the initial plasticity may have represented a maladapted state induced by the perturbation of a cold environment. Therefore, subsequent adaptation to cold will restore the expression/splicing back to the ancestral optimal level. On the other hand, some of the initial plasticity may be beneficial in the short-term. The naïve populations may respond to a new environment by altering expression/splicing immediately, e.g., stress response, which allows them to persist in the environment with certain costs. Once better mechanisms of coping with the environment have evolved, the stress response may no longer be induced. Without stress response activation, regulation for the cold-adapted population in the cold environment could revert toward that observed for the warm-adapted population in the warm environment. From the GO enrichment test (Table S2), we do find significant enrichment for GO terms “response to stimulus” and “response to stress” among the genes that show naïve plasticity, although they are not limited to genes showing “neutralizing” changes. The presence of stress response genes among the “neutralizing” category could make sense if warm-adapted flies in the cold environment manifest a stressed transcriptional state, then it could make sense for (less-stressed) cold-adapted flies in that same cold environment to show stress response gene regulation more aligned with that of the warm-adapted population in its home environment. In other cases, novel regulatory states at stress response genes might protect against the challenges of the cold environment.

It is worth noting that, although the neutralizing cases are enriched in *P_ST_* outliers relative to non-outliers, ~20% – 50% of the outliers show “concordant” changes (Fig. 3). For GO enrichment tests for those showing “concordant” plasticity evolution, only the MED pair has significant terms. These significant GO terms in MED are related to the synapse and behavior, which is consistent with other findings that synapse-associated genes are associated with cold tolerance (Mackay et al., 2012; Pool et al. 2017).

Notably, the ETH pair has many more genes/introns showing reversing changes than the other two pairs. The ETH highland population exhibits distinct phenotypic evolution such as darker pigmentation (Bastide et al., 2014), larger body size (Pitchers et al., 2013; Lack et al., 2016b), and reduced reproductive rate (Lack et al., 2016b), raising the possibility that the strong plasticity-reversing expression change in EF may contribute to the unique phenotypic evolution of this high altitude (3050 m) population. We note that the ETH pair was previously found to have unique regulatory evolution in other respects (Huang et al., 2021): a large majority of its adult (and larval) *P_ST_* outliers had increased rather than decreased expression, and it showed a particular importance of trans-regulatory evolution compared with the other population pairs. Hence, while multi-population comparisons like this one may illuminate evolutionary patterns of general significance, the specific ecological and evolutionary context of each population may importantly influence transcriptome-wide patterns of change.

Plasticity can be reduced by environmental canalization (Debat & David, 2001; Liefting et al., 2009), which may or may not share a common basis with genetic canalization (Wagner et al.,1997; Meiklejohn & Hartl, 2002; Flatt 2005; Pesevski & Dworkin 2020). Here we explored whether putatively adaptive regulatory evolution may disrupt genetic canalization, using inbreeding as a broad genetic perturbation. We first observed a transcriptome-wide pattern of warm-adapted populations showing greater vulnerability to the genetic perturbation of inbreeding than cold-adapted populations in all pairs for both expression abundance and intron usage (Fig. 4A&B). One explanation is that a low temperature is a form of environmental stress to the warm-adapted populations but not so much to the cold-adapted ones, and that this stress induces a relatively uniform transcriptomic response. In combination, the environmental perturbation of cold stress may have compromised the ability of warm-adapted organisms to buffer the genetic perturbation of inbreeding and led to a greater reduction in canalization (higher *V_inbred_*/*V_outbred_* ratio) than the less-stressed cold-adapted organisms (Chen et al., 2015). Although this inbreeding-environment interaction has been observed before, the underlying mechanisms are not understood (Kristensen et al., 2006; Reed et al., 2012). Another potential contributor to the apparently greater transcriptome-wide effects of inbreeding on warm-adapted populations is a potentially larger number of recessive deleterious variants. All three of our cold-adapted populations have slightly lower genomic diversity than their warm-adapted counterparts and appear to have gone through mild population bottlenecks after the population pairs diverged (Lack et al. 2016a; Sprengelmeyer et al. 2020). After the initial generations of such a bottleneck, recessive load is expected to be lower in the bottlenecked population than in the non-bottlenecked counterpart (Kirkpatrick and Jarne 2000), and hence our warm-adapted populations would be predicted to harbor greater recessive load than their cold-adapted partners.

Aside from the above transcriptome-wide differences in canalization, adaptation could reduce canalization if selection shifts gene regulation outside of the buffered ancestral range. Previously, evidence of decanalization linked to adaptive evolution in nature was limited to transient developmental stability in insecticide-resistant blowflies (Clarke & McKenzie, 1987; McKenzie & Game, 1987) and wing abnormalities in large-winged Ethiopian *D. melanogaster* (Lack et al. 2016b). Here, we find that genes with putatively adaptive expression changes are more vulnerable to inbreeding in all three population pairs (Fig. 4C; Fig. 4E). These results are consistent with the adaptation-decanalization hypothesis, but they could also be generated if linked deleterious recessive regulatory variants hitchhiked along with favored haplotypes. We reasoned that only the decanalization model predicts that stronger and especially “novel” regulatory changes should result in greater vulnerability to inbreeding, and we found some evidence in support of this prediction. Specifically, we found that (1) particularly for the ETH population pair, outlier genes with novel regulatory changes were significantly more vulnerable to inbreeding, and (2) for the MED and ETH population pairs, neutralizing changes with greater magnitudes showed stronger evidence for decanalization (Fig. S5). It is intriguing that decanalization evidence was strongest for the ETH pair, given that wing size evolution in the cold-adapted member of this pair was the morphological inspiration for the adaptation-decanalization hypothesis (Lack et al. 2016b).

Our findings involving large numbers of putatively adaptive expression traits could hint at a much broader relationship between adaptive evolution and decanalization, at least for gene expression. However, variation in the strength of evidence for decanalization among our population pairs suggests that evolutionary context may be an important factor modulating the relationship between adaptation and canalization. Further work is needed to understand this relationship more clearly. For example, to what extent do adaptive changes by themselves break down canalization, versus decanalization occurring separately as a prerequisite for adaptive trait change? And to the extent that decanalization occurs concurrently with adaptation trait evolution, how often does it occur as a pleiotropic byproduct of the adaptive change itself, versus arising from linked deleterious variants?

In contrast, evidence for decanalization was not observed to a significant degree for putatively adaptive intron usage changes (Fig. 4D). It is possible that our splicing *P_ST_* outliers may be less enriched for targets of strong adaptive evolution than our expression outliers. Alternatively, there may be a meaningful difference in how the two traits are buffered. While our plasticity data suggest that expression and splicing are similarly susceptible to environmental plasticity (Table 1), it is possible that intron usage traits are less vulnerable to the genetic perturbation of inbreeding, perhaps due to differences in the properties of biological networks governing expression and splicing.

In this study, we leveraged parallel adaptive divergence among natural populations of a model organism to broadly examine the relationship between adaptive evolution, thermal environment, and gene regulation. However, much work remains to be done. Our finding that gene expression evolution tends to neutralize ancestral plasticity is consistent with many although not all previous studies (see Introduction), and suggests that many plastic expression changes may be maladaptive in novel environments. For the first time, we also showed a similar pattern for alternative splicing, in the context of intron usage. It will therefore be of interest to assess the degree to which a broader range of traits (gene regulatory and otherwise) show counter-gradient patterns of evolution and plasticity. Our investigation of the relationship between adaptive gene regulatory evolution and decanalization particularly calls out for further research, in light of this topic’s lack of prior study. It was only relatively recently that an instance of morphological evolution in nature was found to yield decanalization of the same structure’s development (wing size in our highland ETH population; Lack et al. 2016b). We find some evidence that putatively adaptive gene expression traits are linked to (genetic) decanalization on a transcriptome-wide scale, which raises the question of whether a more general role for adaptive evolution in undermining canalization may exist. However, there is a considerable need to investigate this relationship across a broader range of organisms and traits.

## Materials and Methods

### RNA sample collection and sequencing

As described in previous publications (Lack et al., 2016a; Pool et al., 2016; Huang et al., 2021), we have three *Drosophila melanogaster* population pairs, each representing cold-adapted and warm-adapted populations from the same region: a Mediterranean pair (France FR and Egypt EG), an Ethiopian pair (EF, EA) and a South African pair (SD, SP). Within each of these six populations, we selected 16 strains and assigned them into eight crosses. These strains had been inbred for eight generations. To reduce within-cross variance, pairs of strains for a cross were chosen based on minimal overlapping heterozygosity genome-wide. Each cross was conducted concurrently at both 25°C, the control warm condition, and 15°C, the derived cold condition (Fig. 1A). Twenty virgin females and 20 males were collected from maternal and paternal lines respectively for each cross and allowed to mate and lay eggs for a week in half-pint bottles. Each bottle contained standard *Drosophila* medium (containing molasses, cornmeal, yeast, agar, and antimicrobial agents). To collect samples for RNA-seq, 30 female F1 offspring were collected 4-5 days after eclosion and shock-frozen in liquid nitrogen immediately. To compare the effects of inbred and outbred on expression variation after adaptive evolution, we also reared four inbred lines at 15°C for each population (Fig. 1A) and collected 30 females 4-5 days after eclosion. 15°C represented the derived environment where the cold-adapted populations underwent selection.

For RNA extraction, 30 females for each sample were homogenized using TissueLyser II (Qiagen, Hilden, Germany). Total mRNA was isolated using the Magnetic mRNA Isolation Kit (New England Biolabs, Ipswich, MA) and cleaned up using RNeasy MinElute Cleanup Kit (Qiagen, Hilden, Germany). Strand-specific libraries were prepared using the NEBNext mRNA Library Prep Reagent Set for Illumina. Libraries were size-selected for approximately 150 bp inserts using AMPureXP beads (Beckman Coulter, CA, USA). The libraries were quantified using Bioanalyzer and manually multiplexed for sequencing. All libraries were sequenced on a HiSeq2500 (V4) with 2 ⍰ 75bp paired-end reads in two flow cells. Numbers of paired-end reads generated for different libraries can be found in Table S1. Data from parental outbred samples, parental inbred samples, and all F1 samples were each largely sequenced on the same flow cell as other samples of the same type, with a minority of samples having initially low depth of coverage subsequently topped off by further sequencing on the same machine.

### Quantifying gene expression and intron usage frequency

The paired-end sequence reads from each of the F1 samples were mapped to the transcribed regions annotated in *D. melanogaster* (release 6, BDGP6.84) using STAR with parameters from ENCODE3’s STAR-RSEM pipeline (Dobin et al., 2013; Li & Dewey, 2011). Numbers of mapped read pairs for different samples can be found in Table S2. For gene expression, the numbers of reads mapped to each gene were quantified using RSEM (Li & Dewey 2011). Reads mapped to the rRNA were excluded in the analysis. Table S3 provides the read counts for each gene in each sample. The expression abundance for each gene was the number of reads mapped to the gene per million reads (standardized by total reads mapped to the transcriptome).

To visually describe the transcriptome variation among samples, we first performed principal component analysis (PCA) for the F1 samples from different temperature conditions across populations. We used DESeq2 (Love et al. 2014) to construct the data object from the matrix of count data output from RSEM and performed variance stabilizing transformation (vst). The top 5000 genes with highest variance across samples at the transformed scale were used for PCA with the function *prcomp* in R.

To quantify intron usage, we used Leafcutter (Li et al., 2018) to estimate the excision frequencies of alternative introns. Leafcutter took the alignment files generated by STAR as input to quantify the usage of each intron. Then Leafcutter formed clusters that contained all overlapping introns that shared a donor or acceptor splice site. The default parameters were used: > 50 reads supporting each intron cluster and < 500kb for intron length. The numbers of intron excision events for different clusters in each sample can be found in Table S3. The intron usage frequency is the number of intron excision events divided by the total events per cluster. It is worth noting that Leafcutter only detects exon-exon junction usage and it is unable to quantify 5’ and 3’ end usage and intron retention, which may be confounded by differential bias among libraries.

### Comparison of naïve plasticity and evolutionary changes

The detailed methods of identifying candidates for adaptive evolution between cold and warm populations can be found in Huang et al. (2021). Briefly, that study used *P_ST_* statistics to quantify gene regulatory divergence (for both expression abundance and intron usage) between cold- and warm-adapted populations in each pair. That approach was used because the goal was to identify transcriptomic traits with unusually high population differentiation relative to within-population variation (representing candidates for local adaptation), as opposed to simply testing whether any significant population difference in expression/splicing existed at all. The upper 5% of *P_ST_* quantile was used to identify outliers that are more likely to have been under adaptive evolution. (evaluated separately for each population pair). While this threshold is necessarily arbitrary, in light of our present study’s focus on transcriptome-wide patterns, it is not essential that every *P_ST_* outlier be a true product of local adaptation. Instead, we simply need a group of genes that is meaningfully enriched for true local adaptation targets, and that contains enough such genes to provide reasonable power for the analyses described below. Given that we are studying population pairs that have occupied contrasting environments for tens of thousands of generations (Sprengelmeyer et al. 2020) and the differ in multiple traits (e.g. Bastide et al. 2016; Lack et al. 2016b; Pool et al. 2017), it seems reasonable to propose that *P_ST_* outliers are enriched for targets of local adaptation, and that differences between outliers and non-outliers are informative regarding the characteristics of genes under adaptive transcriptomic evolution. However, we note that factors such as population differences in organ size could also influence our whole-organism RNA-seq data, as further discussed in Huang et al. 2021.

To quantify plasticity, we used the difference in gene expression between rearing conditions for each cross. For expression abundance, we calculated the change of expression abundance from cold to warm condition for each of the eight crosses (where expression in both conditions was above 0). To identify genes showing naïve plasticity in warm-adapted populations, we required consistent expression changes (consistently higher or lower expression for the cross in cold condition than that in warm condition) in at least seven out of eight crosses following the direction of average change among the eight crosses. Similarly for intron usage, we calculated the between-environment differences in intron usage proportion for each of the eight crosses. The criterion to identify plastic splicing in warm-adapted populations was consistent differences in at least seven out of eight crosses showing the same direction as the average difference. In the absence of true plasticity, only 7% of genes should meet this criterion by chance (two-sided binomial calculation). Null simulations in which all 16 values were normally distributed confirmed that the false positive rate should be close to 7%. While a 7% false positive rate is slightly higher than the conventional 5%, it represents a reasonable compromise in light of the data available and the goals of our study. Here, our need is to define a set of plasticity-enriched genes of adequate size to offer power for the analyses described below.

To examine whether adaptation to cold environments tends to enhance or oppose the naïve expression plasticity (Fig. 1B), we first calculated the expression value for every gene in each population at each condition. For expression abundance, the median abundance among eight F1 samples was used as the expression value for a population in a certain condition (*Z*). For intron usage, the corresponding value was the median of intron usage proportion among the eight F1 samples. To determine whether a regulatory trait evolved to enhance or oppose ancestral plastic regulation (Fig. 1C-E), we first examined “naïve plasticity” (warm-adapted population in cold environment, *Z_warm_15°C_*, compared to the warm environment, *Z_warm_25°C_*; Fig. 1B). We then asked whether evolution in the cold-adapted population enhanced or opposed the ancestral naïve plasticity. Formally, we asked whether the differences in trait values between cold- and warm-adapted populations in the cold environment, *E* = *Z_cold_15°C_* – *Z_warm_15°C_*, were in the same direction as the plastic response of the warm population, *P_warm_* = *Z_warm_15°C_* – *Z_warm_25°C_*. If *E* ⍰ *P_warm_* > 0, the evolved regulation was assigned as “concordant” (Fig. 1C). If *E* ⍰ *P_warm_* < 0, the evolved regulation was assigned as “discordant”, and these discordant traits were further partitioned into two categories. We defined as “neutralizing” traits in which the cold population’s “home” value (*Z_cold_15°C_*) evolved to be closer to the warm population’s “home” value (*Z_warm_25°C_*) than predicted by the warm population’s plastic response, and therefore |*E*| < 2|*P_warm_*| (Fig. 1D). In contrast, we defined as “reversing” traits in which an evolved change *E* was both opposing the direction of plasticity *P* and took *Z_cold_15°C_* farther from *Z_warm_25°C_* than predicted by the warm population’s plastic response (|*E*| > 2|*P_warm_*|; Fig. 1E). For both categories where evolved expression was farther from the warm population’s “home” value range (either concordant or reversing), we defined them as “novel”.

The differences in “concordant” to “discordant” ratio between *P_ST_* outliers and non-outliers were tested based on the chi-squared test in R version 3.3.0. for both expression abundance and splicing. It is worth noting that the test assumes that regulation is independent among genes and introns. However, it is possible that the expression or splicing of different genes is causally related because of the shared regulatory network.

Because the calculations of *P_warm_* and *E* included a common value, *Z_warm_15°C_*, any random measurement error on this value may generate an artifact of a negative relationship between *P_warm_* and *E* (Mallard et al. 2018; Ho & Zhang, 2019). We therefore repeated this analysis while subdividing the data for estimating *Z_warm_15°C_*: four random crosses were used to estimate *P_warm_* and the other crosses were used to estimate *E*. We used the latter four crosses of warm-adapted populations at 15°C for measuring *E* and the eight crosses of cold-adapted populations at 15°C to identify *P_ST_* outliers and non-outliers. Then we used the same criteria to categorize the evolutionary changes into the “concordant” or “discordant” and repeated the same analysis for all sets of subdivided data (70 sets). Since we used *Z_warm_15°C_* estimated from different data sets to calculate *P_warm_* and *E*, the potential artifact of a negative relationship between *P_warm_* and *E* should be removed. The fraction of the subdivided data (70 sets) showing an excess of concordant changes in *P_ST_* outliers vs. non-outliers was the p-value for testing whether *P_ST_* outliers had a significantly lower concordant proportion than non-outliers.

### GO enrichment test for expression plasticity

Gene Ontology (GO) enrichment tests were performed using the R package “clusterProfiler” (Yu et al., 2012) based on the fly genome annotation (Carlson 2019). For expression abundance, the gene ID for concordant, neutralizing or reversing outliers were input as the focal gene list while all the genes used in studying expression abundance divergence were input as the universe gene list. For intron usage, they were mapped to gene regions to generate the universe gene list. Genes including concordant, neutralizing or reversing outlier intron junctions were used in the focal gene list. The types of GO terms being tested contained all three sub-ontologies: Biological Process (BP), Cellular Component (CC) and Molecular Function (MF). Selection of over-represented GO terms was based on adjusted p-value < 0.1 using “BH” FDR method (Benjamini & Hochberg, 1995) for each sub-Ontology. Based on the description of the GO terms, only the enriched GO terms with distinct functions were reported. If multiple GO terms were over-represented by the same set of candidate genes, only the term annotated with fewest genes was reported.

### Testing the decanalizing effect of adaptation on gene expression

We examined the level of canalization for gene expression under inbreeding as a genetic perturbation. For each population, we selected four outbred F1 samples from which we also had data from one of the parental inbred lines. For gene expression abundance, we calculated the variance among four inbred samples and that among four outbred samples for each population. For intron usage, we re-ran leafcutter based on all inbred samples (24 samples in total for six populations) and all outbred samples (24 samples). Then we calculated the variance for inbred samples (*V_inbred_*) and for outbred samples (*V_outbred_*) for each population. While variance estimates from only four values are expected to be noisy, our focus is exclusively on transcriptome-wide correlations and not conclusions about individual genes. Only those expression abundance or intron usage traits with variance above 0 for both inbred and outbred samples in both populations for each pair were used in this comparison. We first compared the transcriptome-wide fractions showing *V_inbred_ > V_outbred_* between cold and warm populations for both expression abundance and intron usage. Because the hypothesis is about recent adaptive change to novel states resulting in decanalization, we focused on the set of genes/introns that evolved expression novelty (identified in the previous section). To test the role of selection on canalization, we compared the fractions showing higher *V_inbred_* to *V_outbred_* ratio (*r*) in the cold population (*r_cold_*) than that in the warm population (*r_warm_*) for *P_ST_* outliers with that for non-outlier controls using a chi-squared test. We note that we are only comparing variances between inbred and outbred samples of the same gene from the same population, and not between genes that may have different average expression levels. The comparisons between the *P_ST_* outliers and non-outliers were done for “novelty” category and for “neutralizing” category. For those in the “neutralizing” category, we further classified them into undercompensating (|*E*| < |*P_warm_*|) or overcompensating (|*P_warm_*| < |*E*| < 2|*P_warm_*|) and performed similar analysis for each category.

To distinguish between decanalization model and deleterious model, we performed a generalized linear model (GLM) regression analysis on expression abundance across genes for each population pair:

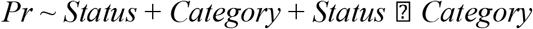

where *Pr* is whether a gene showed *r_cold_* > *r_warm_* as opposed to *r_cold_* > *r_warm_*. *Status* is whether a gene was a *P_ST_* outlier or non-outlier. *Category* is whether a gene was classified as “novelty” or “neutralizing”. We tested for a significant interaction between *Status* and *Category* variables by comparing the above full model against a reduced model without the interaction term. A likelihood ratio test was used to determine significance. We repeated the analysis on genes classified as “neutralizing” to test the effect of overcompensating/undercompensating category on the enrichment of *r_cold_* > *r_warm_* in outliers vs. non-outliers.

## Data Availability

The raw RNAseq reads for samples in the cold condition (15°C) are available from the Sequence Read Archive (SRA) under BioProject PRJNA720479, and those from the warm condition (25°C) are under BioProject PRJNA758705. Table S3 and relevant codes are available at https://github.com/YuhengHuang87/ExpressionPlasticityCanalization

## Acknowledgements

We thank Colin Dewey for helpful discussions. We thank the anonymous reviewers for their helpful comments on the manuscript. This work was funded by NSF DEB grant 1754745 to JEP and by NIH NIGMS grant F32GM106594 to JBL.

