## Supplemental document for "Gene regulatory evolution in cold-adapted fly populations neutralizes plasticity and may undermine genetic canalization"

**Supplementary Figures**

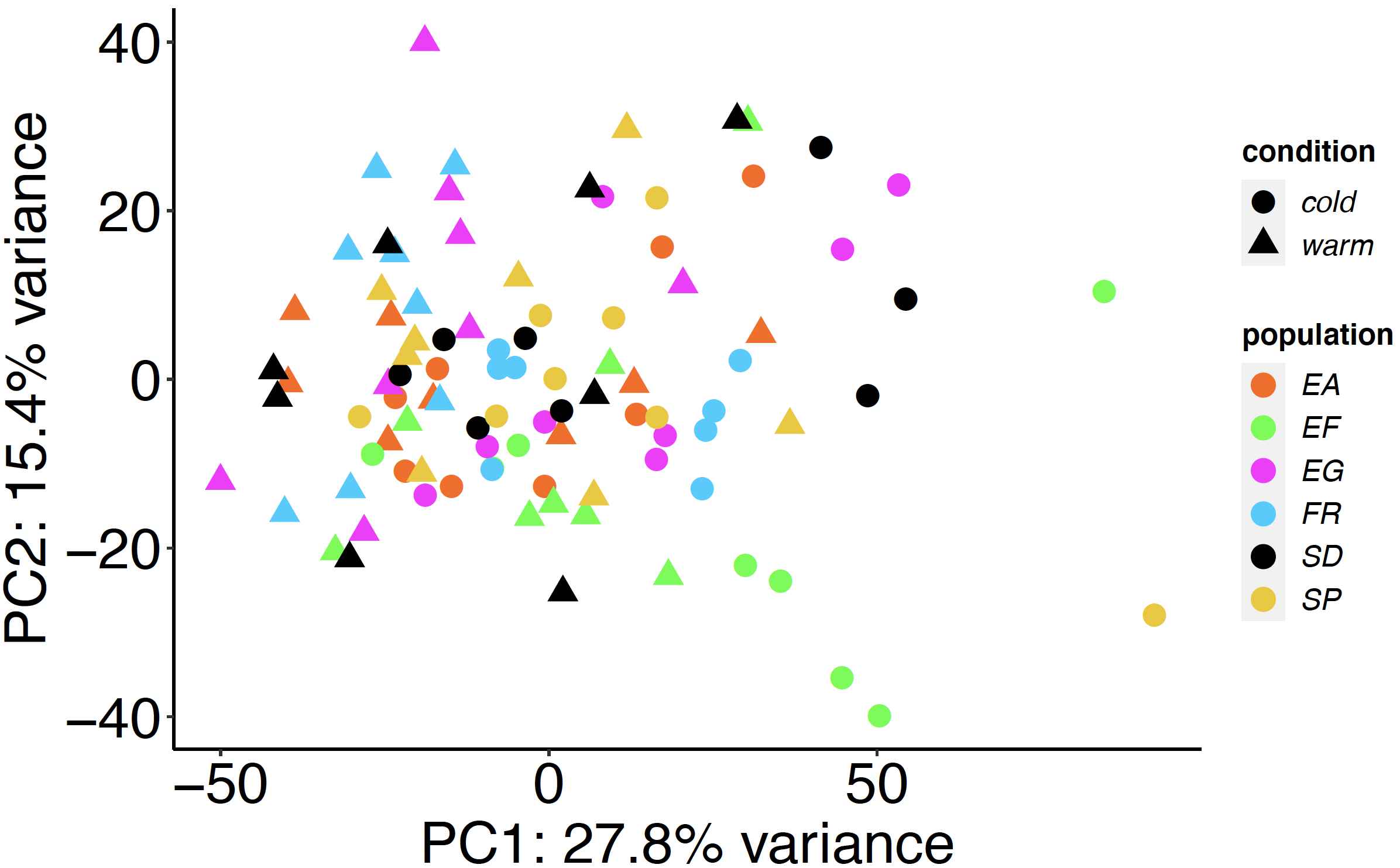

Fig. S1. The PC1 and PC2 for transcriptome abundance among samples from cold and warm environments across populations.

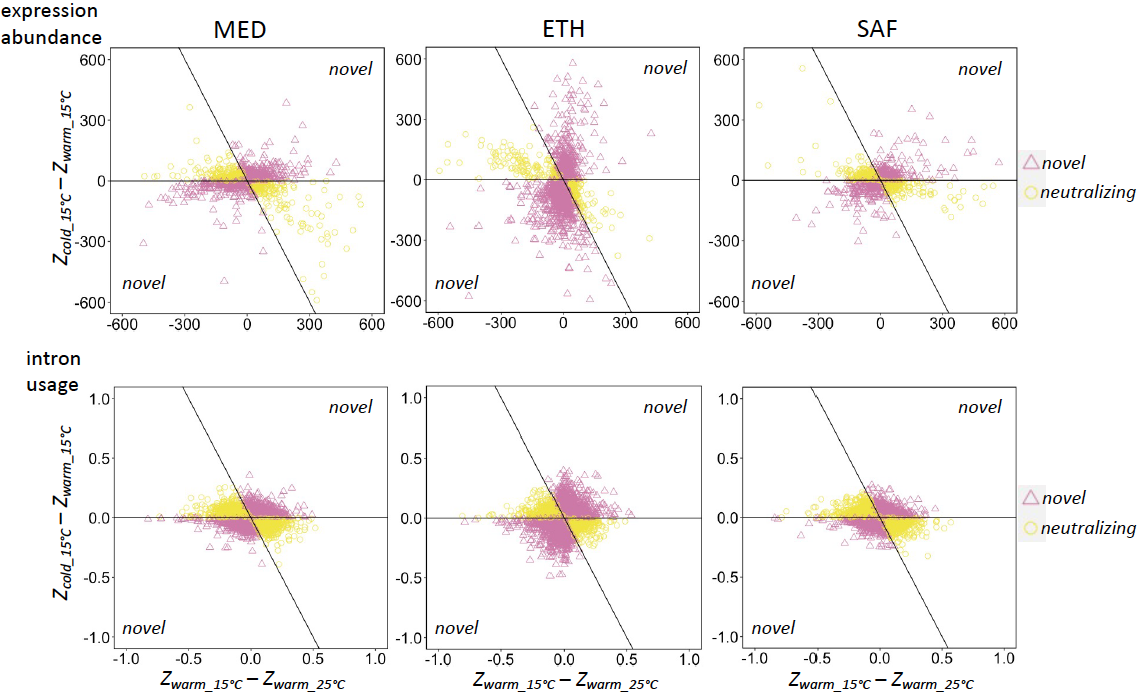
Fig. S2. Gene expression/splicing that is characterized as “novelty” or “neutralizing”. X-axis is the plastic responses between cold and warm environments of the warm population and y-axis is the evolved difference between populations in cold environment for all genes (upper panel) and introns (lower panel).

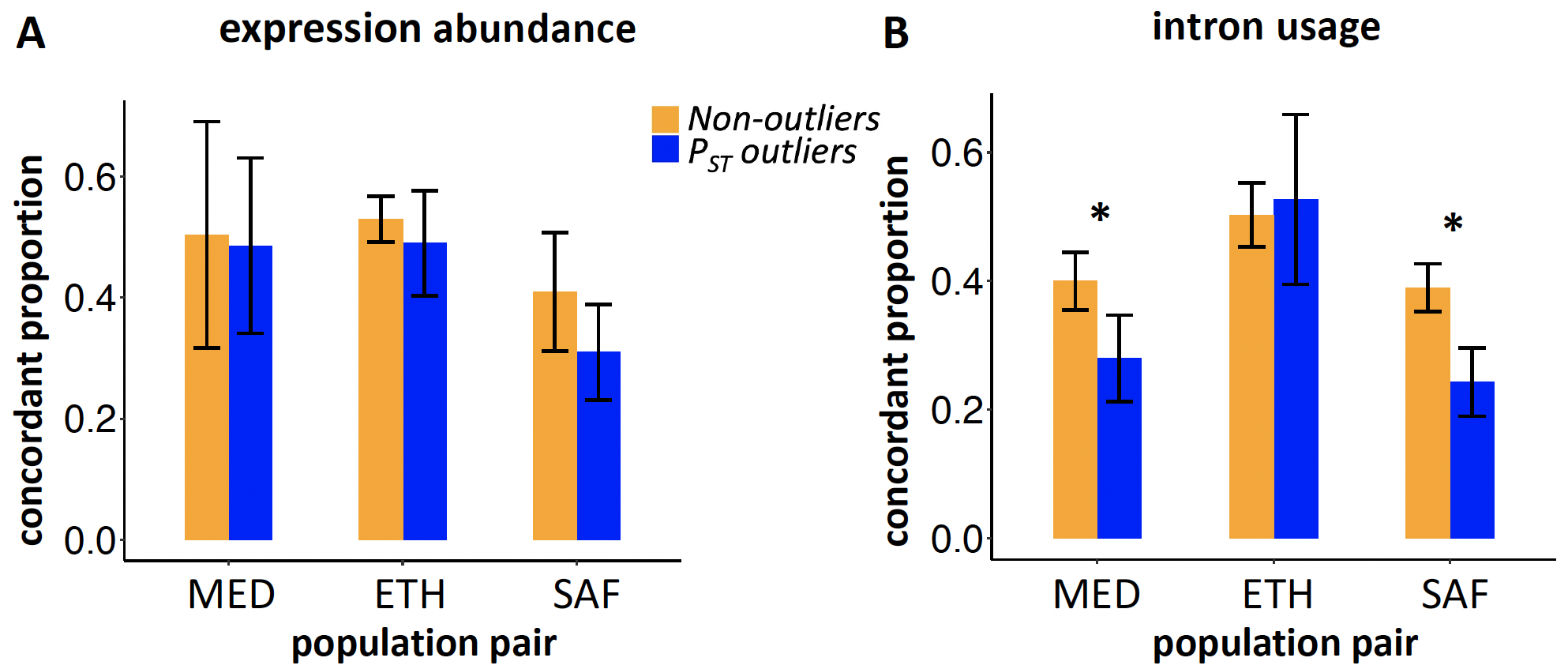

Fig. S3. The proportions of genes/introns classified as “concordant” of naïve plasticity for *P_ST_* outliers and non-outliers. Here the calculations of naïve plasticity value (*P_warm_*) and evolved difference in cold environment (*E*) are based on independent estimations of *Z_warm_15°C_* from two different sets of four crosses. The identification of *P_ST_* outliers are based on the four crosses of warm-adapted populations at 15°C for measuring *E* and the eight crosses of cold-adapted populations at 15°C. There are 70 different combinations of two different sets of four crosses. Means of the concordant proportion for the 70 combinations are plotted. Error bar indicates the standard deviation of the estimated concordant proportion. * indicates the excess of discordant changes in *P_ST_* outliers vs. non-outliers is significant (p < 0.05).

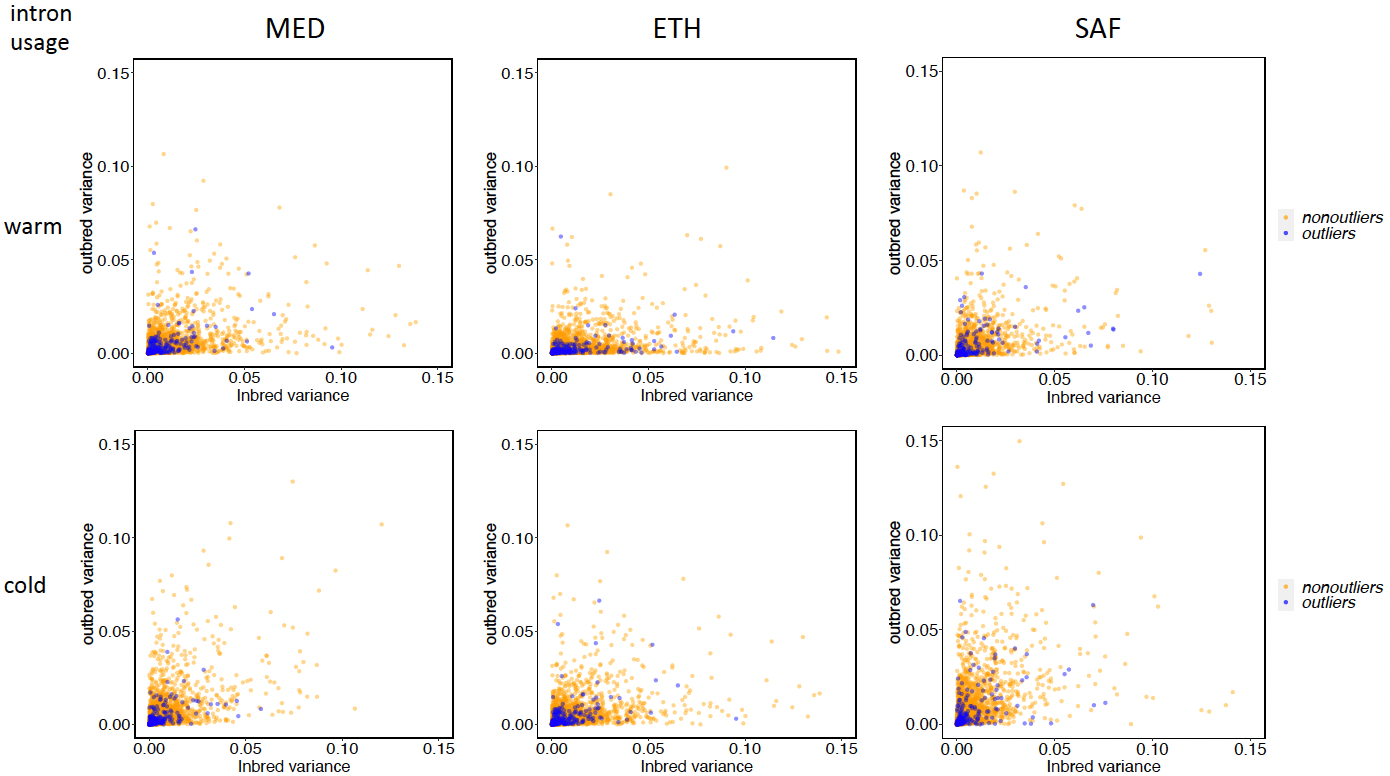

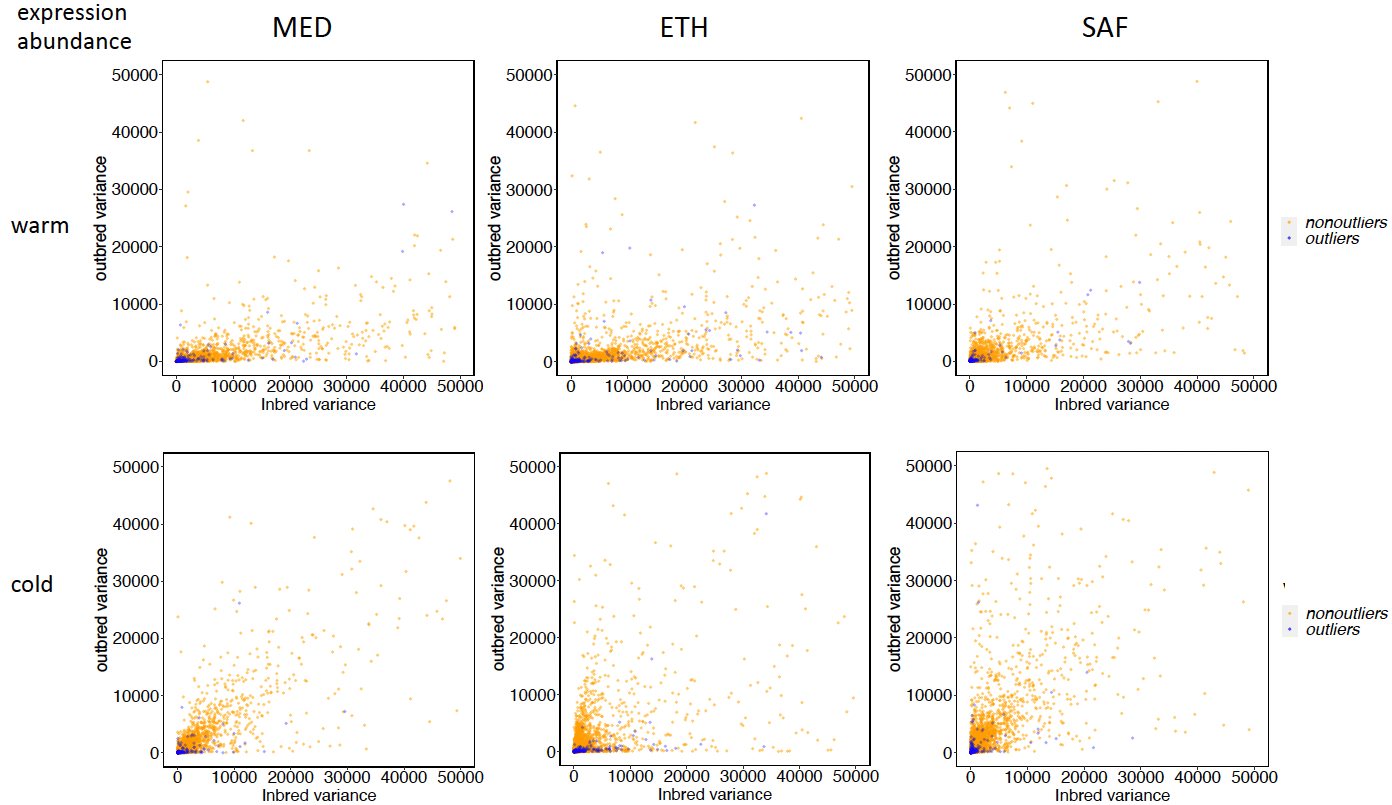

Fig S4. The variance of inbred samples vs. outbred samples for gene expression and intron usage for each population. The *P_ST_* outliers are colored in blue. All the correlations between inbred variance and outbred variance are significant (p < 2.2e-16, spearman correlation).

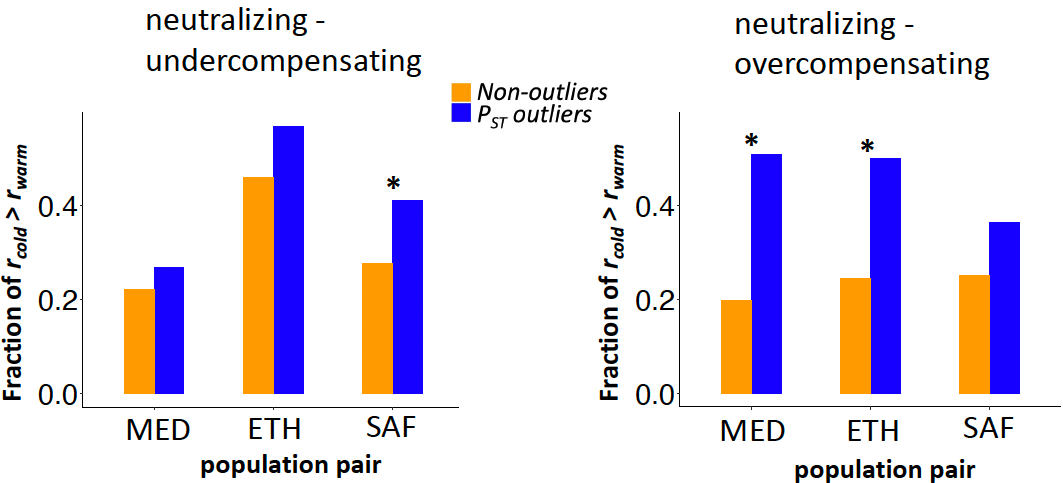

Fig. S5. The fraction of neutralizing expression changes that undercompensate (left) or overcompensate (right) the initial plasticity for which *V_inbred_*/*V_outbred_* (*r*) is higher in a cold population (*r_cold_*) than the respective warm population (*r_warm_*) for outliers and nonoutlier controls. * indicates the proportion is a significant difference between *P_ST_* outliers and non-outliers (p < 0.01).

**Supplementary Tables and Legends**

Table S1. The numbers of reads (75bp paired-end) generated for different samples.

Table S2. The numbers of mapped read pairs for different samples.

Table S3. The read counts for each gene and the intron excision events for each cluster for different samples.

| pair | Mediterranean | Ethiopian | South African |
| --- | --- | --- | --- |
| concordant | 1.cell-cell signaling  2.large ribosomal subunit  3.mitochondrial ribosome  4.neuron part  5.neurotransmitter secretion  6.organellar large ribosomal subunit  7.organellar ribosome  8.plasma membrane receptor complex  9.postsynaptic membrane  10.proteasome core complex  11.receptor complex  12.regulation of neurotransmitter levels  13.unfolded protein binding  14.ribosome  15.spindle  16.spindle microtubule  17.synapse  18.synaptic membrane  19.synaptic signaling | NA | NA |
| neutralizing | 1. amino sugar metabolic process  2. carboxylic acid metabolic process  3. cellular respiration  4. chitin binding  5. chitin metabolic process  6. cuticle development  7. *fatty acid biosynthetic process*  8.glucosamine-containing compound metabolic process  9. monocarboxylic acid biosynthetic process  10. organic acid biosynthetic process  11. organic acid metabolic process  12. oxidoreductase complex  13. oxoacid metabolic process  14. purine nucleoside metabolic process  15. *ribonucleoside metabolic process*  16. *structural constituent of cuticle* | 1. amide biosynthetic process  2. cell projection part  3. cellular amide metabolic process  4. cellular protein metabolic process  5. centrosome organization  6. cytosol  7. cytosolic ribosome  8. intracellular ribonucleoprotein complex  9. large ribosomal subunit  10. long-term memory  11. microtubule cytoskeleton organization  12. microtubule organizing center organization  13. microtubule-based process  14. mitotic spindle organization  15. negative regulation of translation  16. organonitrogen compound biosynthetic process  17. organonitrogen compound metabolic process  18. peptide metabolic process  19. protein metabolic process  20. ribonucleoprotein complex  21. ribosome  22. spindle organization  23. structural molecule activity  24. translation | 1. cell communication  2. cellular response to stimulus  3. DNA binding  4.nucleic acid binding  5. organic cyclic compound binding  6. response to stimulus  7. signal transduction  8. signaling |
| reversing | 1. cell projection | 1. cell cycle checkpoint  2. chromatin  3. chromosomal part  4. chromosome segregation  5. DNA integrity checkpoint  6. DNA metabolic process  7. DNA recombination  8. DNA repair  9. establishment of localization in cell  10. Golgi vesicle transport  hemopoiesis  11. immune effector process  12. immune system process  13. intracellular signal transduction  14. macromolecule catabolic process  15. male meiosis  16. negative regulation of biological process  17. negative regulation of cell cycle  18. nuclear chromosome part  19. nuclear division  20. nucleic acid metabolic process  21. organelle fission  22. organic cyclic compound metabolic process  23. organic substance transport  24. protein complex  25. regulation of catabolic process  26. regulation of macromolecule metabolic process  27. regulation of metabolic process  28. response to extracellular stimulus  29. response to starvation  30. response to stress  31. ribosome biogenesis | NA |

Intron usage

| pair | Mediterranean | Ethiopian | South African |
| --- | --- | --- | --- |
| concordant | 1. behavior  2. biosynthetic process  3. cellular biosynthetic process  4. cognition  5. courtship behavior  6. embryo development  7. establishment of synaptic vesicle localization  8. establishment of vesicle localization  9. learning or memory  10. organelle localization  11. organic substance biosynthetic process  12. regulation of metabolic process  13. response to stimulus  14. synaptic vesicle transport  15. system development  16. vesicle-mediated transport | NA | 1. protein binding |
| neutralizing | NA | NA | NA |
| reversing | NA | NA | 1. cell fate commitment |

Table S4. GO enrichment for the concordant, neutralizing and reversing for expression abundance and intron usage across transcriptomes. GO terms in italics are shared with the GO terms for population differentiation outliers at the adult stage identified in Huang et al. 2021.
